# New insights of zoogeographical distribution of Himalayan goral (*Naemorhedus goral*) from Indian Himalayan Region

**DOI:** 10.1101/2020.08.05.239087

**Authors:** Bheem Dutt Joshi, Vinay Kumar Singh, Hemant Singh, Ashutosh Singh, Sujeet Kumar Singh, Kailash Chandra, Lalit Kumar Sharma, Mukesh Thakur

## Abstract

In the present study, we explored the intraspecific genetic variation and phylogeographic relationship among all the reported species in the genus *Naemorhedus* distributed in a wide range of habitats. The Bayesian based phylogeny demonstrated that Himalayan goral, is a highly diverged species from the other reported species of gorals. We claim the presence of two valid sub-species of Himalayan goral, i.e. *N. g. bedfordi* and *N. g. goral*, distributed in the western and central Himalaya, respectively. The comparative analysis with the inclusion of data available from different ranges, suggests the presence of plausibly six species of gorals across the distribution with a few valid subspecies. Further, we report that *N. griseus* is a valid species and not the synonyms of *N. goral* considering the observed discrepancy in the available sequences. We recommend all the sub-species present at distant locations may be considered as Evolutionary Significant Units (ESUs) and, therefore, appeal to provide them special attention for long term conservation and management.

## 1. INTRODUCTION

Gorals that inhabit in varying elevation (Johnsingh 1992, Hoffmann et al. 2008, Srivastava & Kumar 2018) and distributed in a wide range of habitats (Valdez 2011, Xiong et al. 2013), encompass ambiguity in their species taxonomy (Grubb 2005, Mori et al. 2019). Interestingly, number of species in the genus *Naemorhedus* has been evaluated time to time but still, there is no single consensus on the number of goral species exist (Shukla et al. 2018, Mori et al. 2019, Li et al. 2020). In this context, Valdez (2011) reported six species exist under the genus *Naemorhedus.* However, the International Union for Conservation of Nature (IUCN) still recognizes only the four species i.e. Himalayan goral (*Naemorhedus goral*), Chinese goral (*N. caudatus* and *N. griseus*), and Red goral (*N. bailey*) (Duckworth & MacKinnon 2008, Duckworth et al. 2008a b). Three species of gorals i.e. *N. caudatus, N. griseus* and *N. bailey* are listed as *‘Vulnerable’* whereas Himalayan goral, *N. goral* is classified as ‘*Near Threatened*’ under the IUCN red list category (Duckworth & MacKinnon 2008). All four species of gorals are categorised under the *Appendix I* of CITES and illegal trade of any parts/ products of the listed species is prohibited (Hemley 1994, Wijnstekers 2011). Himalayan gorals possess two sub-species and known to be distributed in all along the Himalayan region of India, Pakistan, Bhutan and Nepal (Grubb 2005, Thapa et al. 2011, Valdez 2011, Shukla et al. 2018). While a few studies are available on the status, distribution, feeding observations and ecological requirements of the Himalayan goral (Ilyas & Khan 2003, Fakhar-i-Abbas et al. 2008, Dar et al. 2012, Srivastava & Kumar 2018), but phylogeographic patterns and inter/intra-specific genetic variations have not been well explored except (Guha et al. 2006, Shukla et al. 2018). However, goral populations in Himalayas are threatened by experiencing habitat fragmentation, degradation, changes in land use patterns, human disturbances, and illegal poaching (Yang et al. 2013, Challender et al. 2015). Further, the existence of the number of species has always been questioned with several rounds of reassignment of six species into three species (Mori et al. 2019) and then to five species (Li et al. 2020). We observed in the current classification of goral species drawn by the phylogenetic species concept, samples from the IHR have not been included. Though, Valdez (2011) suggested the presence of two species, N. *bedfordi* in the western Himalaya and *N. goral* in central and eastern Himalaya. Therefore, we aimed this study to unveil the species composition of Himalayan goral inhabiting in IHR and undertake a comparative assessment with the other goral species.

## 2. MATERIALS AND METHODS

### 2.1. Sampling, PCR and sequencing

We collected a total 60 faecal pellets of Himalayan goral, (#40 from the Uttarkahsi district of the Uttarakhand State in Western Himalayas and #20 from the East Sikkim district of the Sikkim State in the Central Himalayas during the 2018-2019 (Table S1). All the samples were air dried after collection and stored in silica and processed for the DNA extraction using QIAamp DNA Stool Mini kit (Qiagen, Germany). The partial fragment of mitochondrial cytochrome b gene was amplified with all DNA extracts using the universal primers following (Verma & Singh 2003) in a 20 μl reaction volume containing 3.0 μl DNA template of 30–50 ng, 1.5 mM MgCl_2_, 2x PCR Buffer, 0.5 unit Taq DNA polymerase, 50μM dNTPs and 0.5 μM of each primer. PCR thermal cycle was as follows: an initial denaturation for 5 minutes at 94 °C, followed by 40 cycles of denaturation for 45 Sec at 94 °C, annealing at 55 °C for 60 Sec. The cycle sequencing PCR were carried out using the BigDye™ Terminator v3.1 Cycle Sequencing Kit (Applied Biosystems, USA) and cleaned products were sequenced on the ABI 3730 Genetic Analyzer (Applied Biosystems, USA).

### 2.2. Data Analysis

#### Sequence validation, quality checks and data mining

Raw sequences were validated using Sequencher 4.7 (Gene Codes Corporation, USA). Sequences that have low Q values (<30) were not processed for further analysis. All available sequences of different species of goral were retrieved from NCBI (http://www.ncbi.nlm.nih.gov) for comparative phylogenetic assessment (Table S1). Further, pairwise sequence divergence was calculated using the Kimura 2 parameters (K2P) distance matrix in the MEGA X (Kumar et al. 2018).

#### Phylogenetic reconstruction

Bayesian-based phylogeny was reconstructed using BEAUti v 1.6.1 and BEAST v.1.10.4 (Suchard et al. 2018). We applied the best fit model HKY selected by Model test 3.6 (Nylander 2004) with the AIC and BIC criteria. We performed the Markov Chain Monte Carlo (MCMC) runs for 20 million generations sampled at every 1000^th^ tree, where the first 10% of trees sampled were treated as burn. Tree-Annotator v1.8.1 was used to select the maximum clade credibility (MCC) tree (Bouckaert et al. 2014). The MCC tree was visualized in the FigTree v.1.3.1 (Rambaut 2009).

## 3. RESULTS AND DISCUSSION

Sequences generated in the present study well covered the distribution of Himalayan goral in the western (#26 from Uttarkashi district of Uttarakhand State) and central (#6 east Sikkim district of Sikkim State) Himalayas. Thirty-two sequences yielded seven haplotypes from the western and two haplotypes from the central Himalayas. All the nine novel haplotypes are submitted to GenBank (MT845345 - MT845353) We obtained intra-specific sequence divergence between western and central Himalaya was 0.003–0.030 (Table S2). However, comparative analysis by inclusion of goral sequences from Himalaya and other range countries like China, Myanmar, Thailand yielded inter species sequence divergence from 0.03–0.16 (Table S3). Interestingly, Bayesian based phylogeny grouped all sequences into six clades i.e. clade 1 and 2 represented Chinese gorals (*N. caudatus* and *N. griseus*) with some ambiguous sequences of the Himalayan goral (*N. goral*), clade 3, 4 and 5 represented red gorals (*N. bailey* and *N. cranbrooki* and *N. evansi*) and clade 6 represented the Himalayan goral (*N. goral*). The Himalayan goral clade 6 further contained two sub-clades, one each originated from central and western Himalayas (subclade 6a and 6b; Fig. 1). The sequences generated in the present study formed the basal clade that represented two sub-species (*N. g. bedfordi* and *N. g. goral*) which were genetically most divergent lineage from the other reported goral species (Fig. 1). Further, clade 2 contained sequences of both species of Chinese (*N. griseus)* and Himalayan goral *(N. goral).* While, Mori et al. (2019) and Li et al. (2020) suggested that *N. griseus* is the synonyms of the *N. goral.* However, our results were in disagreement with the assignment of Chinese goral-*N. griseus* with *N. goral.* Instead, we raised concern on the labeling of the sequences as *N. goral* that clustered in clade-2 along with *N griseus.* Plausibly, these samples might have been collected either from the distribution range of *N griseus* or if originated from captive facilities then they were mis-identified as *N. goral.* In this context, one of the sequence with Genbank ID MG865962 named as *N. goral* was originally the complete mitochondrial genome of the *N. caudatus griseus* (Yao et al. 2019). Similarly, sequence with Genbank ID MG591488 sampled at the Wutai county in Shanxi Province, China (Ren et al. 2018) and sequence with Genbank ID KP787820 collected at the Tangjiahe Natural Reserve, Sichuan Province, China (Liu & Jiang 2017) were localities fall under the distribution range of *N. griseus*. So, we caution the use of these submission in drawing phylogenetic inferences. Further, one sequence collected from the Mount Qomolangma (Everest) Nature Reserve in Tibet Province exceptionally clustered with clade 2 (Yang et al. 2013). This sequence also raised concern and require further confirmation. In the present study, we made available the competent data of Himalayan goral, *N. goral* from IHR and demonstrated that Himalayan gorals are distinct from all other species of gorals (Fig. 1). The results further evident to prove that goral distributed in India, both in the western and central Himalaya is Himalayan goral (*Naemorhedus goral*; Groves & Grubb 1985) with two valid sub-species *i.e. N. g. bedfordi* and *N. g. goral.* However, Velzen (2011) reported *N. g. bedfordi* as a distinct species (*N. bedfordi*), distributed in the western Himalaya. However, we supported to classify this as a sub-species of *N. goral* due to relatively low sequence divergence (0.012-0.030). Further, our observations also corroborated with an earlier study that reported Indian goral sequences showed a high sequence divergence and formed divergent clade with all other gorals (Shukla et al. 2018).

**Figure 1.**
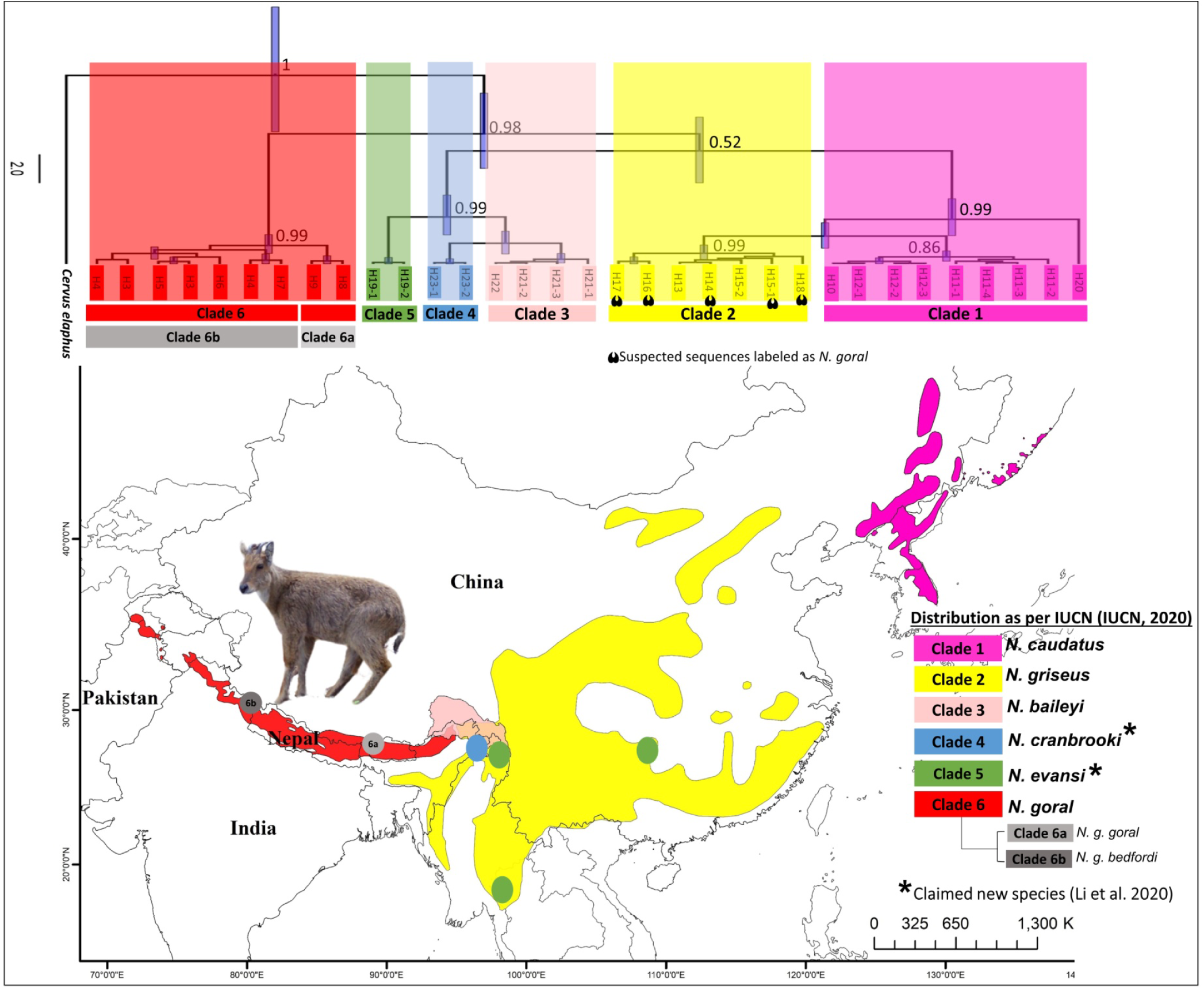
Phylogenetic relationship of different goral species using the partial fragment of mitochondrial cytochrome b gene. The four colors represent the distribution ranges of four goral species as per IUCN (Accessed June 2020: Duckworth, & MacKinnon, 2008; Duckworth et al. 2008a Duckworth et al., 2008b) and two addition dots of sky blue and green represents the distribution of two newly claimed species by Velzen, (2011) and Li et al. (2020) from the Northern Myanmar and Southern China. Above the clades posterior probability are shown. *Cervus elaphus* (NC_007704.2) was used as outgroups for analysis.

Hence, the major inferences from the present study are (1). Indian Himalayan region encompasses a single species of goral which is diverged into two distinct subspecies present in the western and central Himalayas (2). There are potential ambiguities of the sequences available of Himalayan goral, *N. goral* from the China; and *N. griseus* should not be considered as the synonyms of *N. goral.* Alternatively, *N. griseus* may be considered as a distinct different species following Velzen, (2011) and we questioned the existence of *N. goral* in that region. (3). We support the presence of six species (*N. caudatus, N. griseus, N. goral, N. bailey, N. cranbrooki and N. evansi*) which are then divided into subspecies level (Groves & Grubb 1985, Li et al. 2020). All the sub-species present at distant locations should be monitored considering important Evolutionary Significant Units (ESUs), which are evolving and require special management attention and may seek taxonomically up-gradation in future. We also suggest the extensive sampling of goral species from entire distribution ranges and generation of high depth sequencing data to delineate the ecological/ or geographic boundaries and also to understand their sympatric coexistence in different habitats due to increasing number of species.

## Acknowledgements

Authors thanks to the Principal Chief Conservator of Forest (PCCF) and Chief Wildlife Warden (CWLW), of Uttarakhand and Sikkim for granting the necessary permission to carry out research work. We also acknowledge the researcher of NMHS Large Grant project of ZSI who have helped in the samples collection. The study was supported by the funds received from National Mission for Himalayan Studies, Ministry of Environment, Forest and Climate Change (MoEF&CC), New Delhi, India (Grant No. NMHS/2017-18/LG09/02/476).

## AUTHORS’ CONTRIBUTION

BDJ undertaken field survey and collected samples, VKS Data analysis, MT BDJ conceptualized the idea and edited the MS. BDJ, SKS, LKS, and MT finalized the manuscript. LKS and MT coordinated the project funded under the National Mission Himalayan Studies (NMHS) of Ministry of Environment, Forest and Climate Change (MoEF&CC) and KC supervised the overall activities and provided all the logistic support and administrative approval.

