## Supplementary files for "New insights of zoogeographical distribution of Himalayan goral (*Naemorhedus goral*) from Indian Himalayan Region"

Table S1. Details of sequences used in the present study.

| **Accession No.** | **Species** | **Location** |
| --- | --- | --- |
| MT845345 | *Naemorhedus goral* | Uttarakhand, India |
| MT845346 | *Naemorhedus goral* | Uttarakhand, India |
| MT845347 | *Naemorhedus goral* | Uttarakhand, India |
| MT845348 | *Naemorhedus goral* | Uttarakhand, India |
| MT845349 | *Naemorhedus goral* | Uttarakhand, India |
| MT845350 | *Naemorhedus goral* | Uttarakhand, India |
| MT845351 | *Naemorhedus goral* | Uttarakhand, India |
| MT845352 | *Naemorhedus goral* | Sikkim, India |
| MT845353 | *Naemorhedus goral* | Sikkim, India |
| FJ469673.1 | *Naemorhedus caudatus* | Unknown |
| EU259117.1 | *Naemorhedus caudatus* | Everland Zoo, South Korea |
| AY356357.1 | *Naemorhedus caudatus* | Unknown |
| EU259115.1 | *Naemorhedus caudatus* | Unknown |
| EU259114.1 | *Naemorhedus caudatus* | Unknown |
| EU259108.1 | *Naemorhedus caudatus* | Everland Zoo, South Korea |
| EU259116.1 | *Naemorhedus caudatus* | Everland Zoo, South Korea |
| EU259113.1 | *Naemorhedus caudatus* | Everland Zoo, South Korea |
| FJ207532.1 | *Naemorhedus griseus* | MNHN Museum Samples, France |
| MG591488.1 | *Naemorhedus griseus/goral* | Wutai County in Shanxi Province, China |
| JX188255.1 | *Naemorhedus griseus/goral* | Averaes National Park, Tibbet, China |
| KF500173.1 | *Naemorhedus griseus/goral* | Anhui, China |
| MG865962.1 | *Naemorhedus griseus/goral* | Western Sichua, China |
| KT878720.1 | *Naemorhedus griseus/goral* | Tangjiahe Natural Reserve, Sichuan Province, China |
| EU259118.1 | *Naemorhedus griseus/goral* | Singapore Zoo, Singapore |
| MF155891.1 | *Naemorhedus griseus/evansi* | Guizhou’s Fanjingshan National Nature Reserve, China. |
| JN632664.1 | *Naemorhedus griseus/evansi* | Thailand |
| U17861.1 | *Naemorhedus caudatus* | San Diego Zoo, USA |
| KP203894.1 | *Naemorhedus baileyi* | Shanghai Zoo, China |
| JN632663.1 | *Naemorhedus baileyi* | Rotterdam Zoo, Netherlands |
| JX506309.1 | *Naemorhedus baileyi* | Shanghai Zoo, China |
| JX506310.1 | *Naemorhedus baileyi* | Shanghai Zoo, China |
| MN853098.1 | *Naemorhedus cranbrooki* | Putao County, Northern Myanmar |
| MN853099.1 | *Naemorhedus cranbrooki* | Putao County, Northern Myanmar |

Table S2- Pairwise sequence divergence of different haplotype of gorals using the partial fragment of mitochondrial cytochrome b gene.

|  | H1 | H2 | H3 | H4 | H5 | H6 | H7 | H8 | H9 | H10 | H11 | H12 | H13 | H14 | H15 | H16 | H17 | H18 | H19 | H20 | H21 | H22 | H23 |
| --- | --- | --- | --- | --- | --- | --- | --- | --- | --- | --- | --- | --- | --- | --- | --- | --- | --- | --- | --- | --- | --- | --- | --- |
| H1 |  |  |  |  |  |  |  |  |  |  |  |  |  |  |  |  |  |  |  |  |  |  |  |
| H2 | 0.003 |  |  |  |  |  |  |  |  |  |  |  |  |  |  |  |  |  |  |  |  |  |  |
| H3 | 0.009 | 0.012 |  |  |  |  |  |  |  |  |  |  |  |  |  |  |  |  |  |  |  |  |  |
| H4 | 0.006 | 0.009 | 0.003 |  |  |  |  |  |  |  |  |  |  |  |  |  |  |  |  |  |  |  |  |
| H5 | 0.015 | 0.018 | 0.006 | 0.009 |  |  |  |  |  |  |  |  |  |  |  |  |  |  |  |  |  |  |  |
| H6 | 0.012 | 0.015 | 0.003 | 0.006 | 0.003 |  |  |  |  |  |  |  |  |  |  |  |  |  |  |  |  |  |  |
| H7 | 0.018 | 0.021 | 0.015 | 0.012 | 0.015 | 0.012 |  |  |  |  |  |  |  |  |  |  |  |  |  |  |  |  |  |
| H8 | 0.015 | 0.018 | 0.012 | 0.015 | 0.018 | 0.015 | 0.027 |  |  |  |  |  |  |  |  |  |  |  |  |  |  |  |  |
| H9 | 0.018 | 0.021 | 0.015 | 0.018 | 0.021 | 0.018 | 0.030 | 0.003 |  |  |  |  |  |  |  |  |  |  |  |  |  |  |  |
| H10 | 0.153 | 0.148 | 0.157 | 0.152 | 0.157 | 0.153 | 0.161 | 0.157 | 0.161 |  |  |  |  |  |  |  |  |  |  |  |  |  |  |
| H11 | 0.153 | 0.148 | 0.157 | 0.152 | 0.157 | 0.153 | 0.161 | 0.157 | 0.161 | 0.006 |  |  |  |  |  |  |  |  |  |  |  |  |  |
| H12 | 0.148 | 0.144 | 0.153 | 0.148 | 0.153 | 0.148 | 0.157 | 0.153 | 0.157 | 0.003 | 0.003 |  |  |  |  |  |  |  |  |  |  |  |  |
| H13 | 0.153 | 0.149 | 0.149 | 0.144 | 0.149 | 0.145 | 0.153 | 0.157 | 0.161 | 0.036 | 0.036 | 0.033 |  |  |  |  |  |  |  |  |  |  |  |
| H14 | 0.144 | 0.148 | 0.141 | 0.136 | 0.141 | 0.137 | 0.145 | 0.149 | 0.153 | 0.036 | 0.036 | 0.033 | 0.006 |  |  |  |  |  |  |  |  |  |  |
| H15 | 0.148 | 0.144 | 0.144 | 0.140 | 0.145 | 0.141 | 0.149 | 0.153 | 0.157 | 0.033 | 0.033 | 0.030 | 0.003 | 0.003 |  |  |  |  |  |  |  |  |  |
| H16 | 0.153 | 0.148 | 0.148 | 0.144 | 0.149 | 0.144 | 0.153 | 0.157 | 0.161 | 0.036 | 0.036 | 0.033 | 0.006 | 0.006 | 0.003 |  |  |  |  |  |  |  |  |
| H17 | 0.148 | 0.144 | 0.144 | 0.140 | 0.145 | 0.141 | 0.149 | 0.153 | 0.157 | 0.039 | 0.039 | 0.036 | 0.009 | 0.009 | 0.006 | 0.003 |  |  |  |  |  |  |  |
| H18 | 0.153 | 0.148 | 0.148 | 0.144 | 0.149 | 0.144 | 0.153 | 0.157 | 0.161 | 0.036 | 0.036 | 0.033 | 0.006 | 0.006 | 0.003 | 0.006 | 0.009 |  |  |  |  |  |  |
| H19 | 0.125 | 0.121 | 0.129 | 0.125 | 0.137 | 0.133 | 0.141 | 0.129 | 0.133 | 0.128 | 0.128 | 0.124 | 0.128 | 0.128 | 0.124 | 0.120 | 0.124 | 0.120 |  |  |  |  |  |
| H20 | 0.149 | 0.145 | 0.153 | 0.149 | 0.162 | 0.157 | 0.166 | 0.153 | 0.157 | 0.069 | 0.069 | 0.066 | 0.037 | 0.043 | 0.040 | 0.043 | 0.046 | 0.043 | 0.141 |  |  |  |  |
| H21 | 0.128 | 0.124 | 0.140 | 0.136 | 0.140 | 0.136 | 0.144 | 0.140 | 0.144 | 0.123 | 0.115 | 0.119 | 0.130 | 0.130 | 0.126 | 0.123 | 0.126 | 0.123 | 0.056 | 0.164 |  |  |  |
| H22 | 0.124 | 0.120 | 0.136 | 0.132 | 0.136 | 0.132 | 0.140 | 0.136 | 0.140 | 0.119 | 0.112 | 0.115 | 0.127 | 0.126 | 0.123 | 0.119 | 0.123 | 0.119 | 0.059 | 0.160 | 0.003 |  |  |
| H23 | 0.140 | 0.136 | 0.152 | 0.148 | 0.152 | 0.148 | 0.156 | 0.152 | 0.156 | 0.119 | 0.119 | 0.115 | 0.127 | 0.126 | 0.123 | 0.119 | 0.115 | 0.119 | 0.059 | 0.168 | 0.027 | 0.030 |  |

H1-H7; *N. goral*; H10-H12, H20*, N. caudatus*; H13-H18, *N. griseus*, H21-22, *N. baileyi*; H19, *N. evansi;* H23 *N. cranbrooki*

Table S3. Pairwise sequence divergence of different species of gorals using the partial fragment of mitochondrial cytochrome b gene.

|  | *N. caudatus* | *N. griseus* | *N. baileyi* | *N. cranbrooki* | *N. evansi* | *N. goral* |
| --- | --- | --- | --- | --- | --- | --- |
| *N. caudatus* |  |  |  |  |  |  |
| *N. griseus* | 0.038 |  |  |  |  |  |
| *N. baileyi* | 0.095 | 0.086 |  |  |  |  |
| *N. cranbrooki* | 0.096 | 0.084 | 0.021 |  |  |  |
| *N. evansi* | 0.128 | 0.128 | 0.164 | 0.059 |  |  |
| *N. goral* | 0.113 | 0.107 | 0.100 | 0.110 | 0.125 | 0.00 |
